# Identification of snails and parasites of medical importance via convolutional neural network: an application for human schistosomiasis

**DOI:** 10.1101/713727

**Authors:** Zac Yung-Chun Liu, Andy J. Chamberlin, Pretom Shome, Isabel J. Jones, Gilles Riveau, Raphael A. Ndione, Lydie Bandagny, Nicolas Jouanard, Paul Van Eck, Ton Ngo, Susanne H. Sokolow, Giulio A. De Leo

**Author notes:** Send correspodence to and.

## Abstract

Schistosomiasis is a debilitating parasitic disease infecting over 250 million people with nearly 800 million people at risk worldwide, primarily in sub-Saharan Africa. Transmission to humans involves freshwater snails as intermediate hosts, which are particularly prevalent in developing countries where dams and water resource projects have expanded freshwater snail habitat. At our study sites in the lower Senegal River Basin, we have collected more than 5,500 images of the 7 freshwater snail species (grouped into 4 categories) most frequently encountered in this aquatic ecosystem, 5 of which amplify and transmit either urinary or intestinal human schistosomiasis, with the other 2 species responsible for the transmission of less common parasitic diseases of humans and/or livestock. We have also collected over 5,100 images of 11 classes of trematodes, including human and non-human schistosomes. It takes a great deal of training and expertise to accurately classify these organisms morphologically. In recent years, deep convolutional neural networks (CNNs) have proven to be highly efficient for image recognition tasks across many object categories. Here we demonstrate classification capacity for snail and parasite images and test our model’s performance against 8 highly-trained human parasitologists with experience taxonomically classifying snail and parasite species from the Senegal River Basin in West Africa. We establish and train a single CNN end-to-end directly from images with only pixels and labels as inputs. Applying this state-of-the-art algorithm, we are able to classify images of 4 snail categories with 99.64% accuracy and images of 11 parasite categories with 88.14% accuracy, which rivals highly-trained human parasitologists. The trained algorithm could next be deployed to mobile devices for use in remote field settings by local technicians, and significantly improve monitoring snail and parasite occurrence in the field for disease control purposes.

**Author Summary:** Schistosomiasis is a neglected tropical disease (NTD) infecting over 250 million people worldwide. The current approach to mitigate this disease in endemic regions is community or school-based mass drug administration. However, parasites are primarily transmitted through environmental reservoirs where freshwater snails serve as intermediate hosts. People use the contaminated water sources for their daily tasks and get re-infected after drug treatment. Therefore, drug administration alone is not effective for schistosomiasis control in such high transmission settings. Recent studies show that snail population control is essential to reduce disease transmission risks. To discern between parasitic worms of humans in snails and those of other non-human species is a necessary step to precisely quantify transmission risk for human schistosomiasis. However, it takes a great deal of expertise to train lab and field technicians to accurately classify snail and parasite species. Artificial intelligence (AI)-powered computer vision algorithms have recently proven to be highly efficient for image recognition tasks. We have collected thousands of snail and parasite images in Senegal during 2015-2018. Using these images as training data, we developed an AI model that classifies images of 4 snail categories and 11 parasite categories with, in some cases, higher accuracy than well-trained human parasitologists. This model could next be deployed to mobile devices for use in remote field settings to support local technicians to identify transmission hotspots and target control.

## Introduction

Most human parasitic infections, such as schistosomiasis, onchocerciasis, lymphatic filariasis, and malaria, are diseases of poverty: even while effective drugs to treat people exist, the most affected communities are too poor to access them. Global health efforts have attempted to fill this gap by delivering needed medication to rural communities in some of the poorest countries through mass drug administration programs, a strategy that takes a practical approach to control, treating whole communities, affordably, without regard to individual infection status. Yet because many parasites are transmitted through environmental reservoirs, vectors, or intermediate hosts unaffected by this scheme, reinfection after treatment can be a major impediment to achieving control using drugs alone. Thus, complementing human treatment with transmission control, by identifying and eliminating hotspots of vectors and reservoirs in the environment, could be most effective. Yet, this approach is severely hindered by a lack of ecological research capacity in poor countries to identify, monitor, and affordably eliminate relevant vectors and parasites with minimal environmental damage.

Human schistosomiasis is the second most important parasitic disease in the world behind malaria, affecting over 250 million people in over 70 tropical and subtropical countries. The majority of cases occur in sub-Saharan Africa, where, in recent decades, the expansion of dams and water management infrastructure has resulted in the expansion of freshwater snail habitat and human-water contact. These changes have benefited transmission of schistosoma worms, which have a complex life cycle (**Fig 1**) involving: (1) a final vertebrate host (humans), where the adult worms live and reproduce sexually, and (2) an intermediate freshwater snail host where the larval stages of the worms develop via asexual reproduction. People are infected through contact with water contaminated by free-living stages of the parasites that shed daily by infected snails. Upon contact, the parasite penetrates skin and enters the bloodstream. Therefore, infected snails in the freshwater environment are the source of rapid infection and re-infection of humans in endemic areas [4]. And, despite the rise in preventive chemotherapy campaigns in the last several decades, just as many people suffer from schistosomiasis today as did 50 years ago [5].

**Figure 1.**
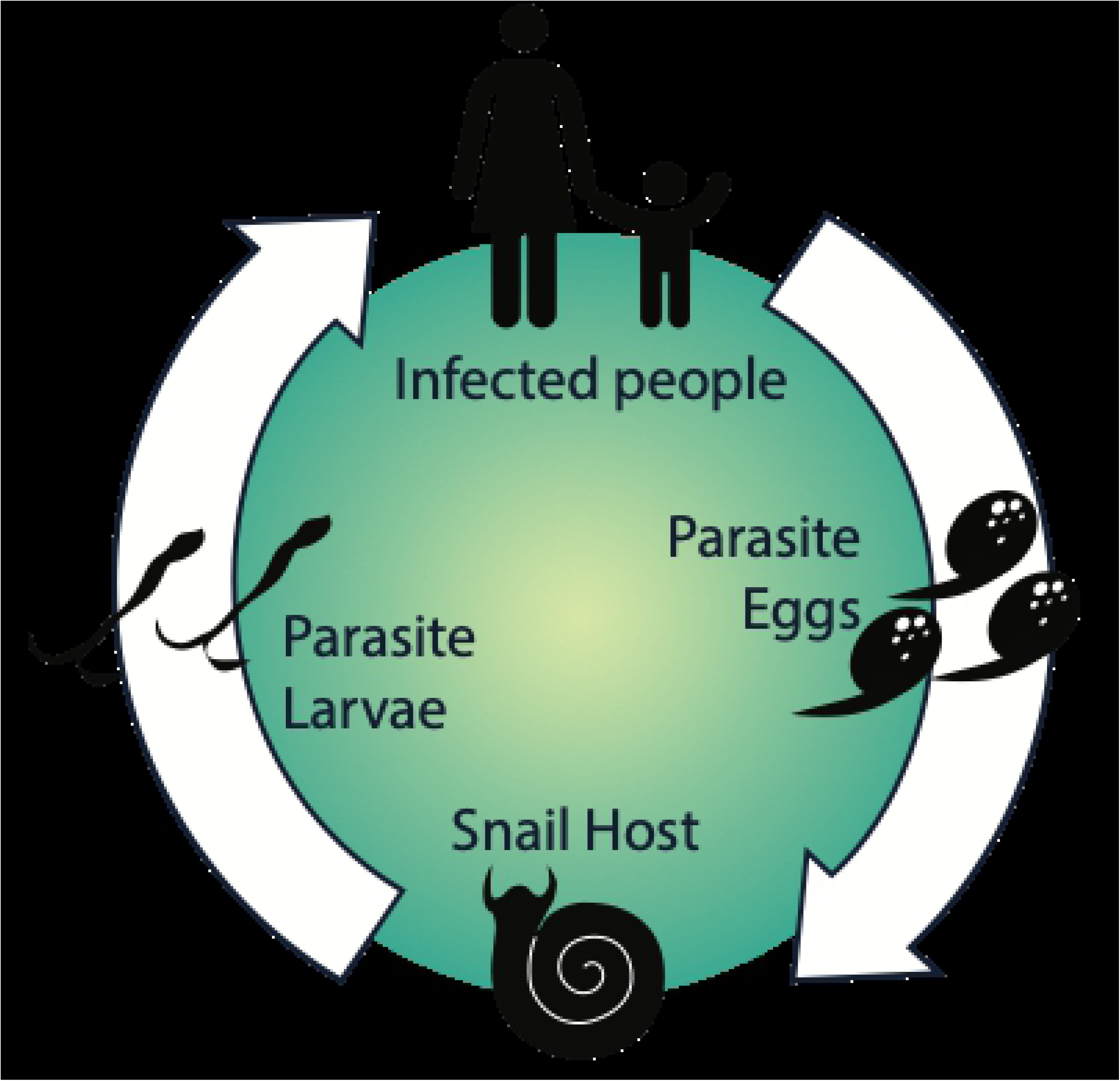
The life cycle of *Schistosoma spp.* parasites. The adult worms live and reproduce sexually within the human host, releasing eggs into the environment that hatch and seek an intermediate freshwater snail host, where the larval stages of the worms develop via asexual reproduction. Cercariae are released from snails and seek human hosts, completing the lifecycle.

Reducing snail populations using molluscicides has been an effective way to reduce transmission of human schistosomiasis because snails serve as intermediate hosts of the parasite [1, 2], but widespread mollusciciding has fallen out of favor because it is expensive and can lead to non-target toxic effects in the environment [3]. New ways to treat environments safely (e.g. natural plant-based molluscicides, or biological control using natural enemies) and to target treatments only where infected snails are abundant are desperately needed.

In sub-Saharan Africa, there are two main species of schistosoma parasites that infect humans and use different snail species as an intermediate host: *Biomphalaria* (for *Schistosoma mansoni*) and *Bulinus* (for *Schistosoma hematobium*) [1]. Detecting the abundance and spatial distribution of both snail species and human schistosoma parasites is essential to design control strategies aimed at reducing the environmental reservoir of the disease and/or administering anthelminthic drugs where transmission risk is highest. Proper classification of snail species and of parasites relevant to human health is the crucial first step to deploy effective control strategies.

One of the most underrated tasks in the fight against schistosomiasis and other helminthiases is accurate classification of snail and parasite species. Attempting this on the basis of morphology is a labor-intensive, time-consuming job that requires a great deal of training experience and expertise. Morphological differences between *Bulinus* spp. and *Biomphalaria* spp. snails at the genus level are clear to any trained naturalist, but distinguishing the specific snail species that are involved in the transmission of human schistosoma parasites might be challenging for students and technicians in their early field and laboratory experience or without continuing training. An additional challenge is that the same snail species that carry human parasites can also be infected with a wide variety of livestock and wildlife parasites, some of which are morphologically similar (and in some cases, even indistinguishable) to human schistosome larvae [6]. As a consequence, visual identification of parasite species during snail dissection can be tricky even for experienced parasitologists. Misidentification of snail infections as human schistosomes, when in fact they are cryptic species that do not harm people, may interfere with cost-effectively deploying badly needed control in areas where resources are limited to begin with.

To address this issue, we developed a computational method with the capacity to automatically identify snails and parasites from photographs taken with a cell phone camera. We apply convolutional neural networks (CNNs), a deep learning application in computer vision, to identify the images and compare results to those of trained human parasitologists.

Holmström et al. (2017) [7] conducted a proof-of-concept study on mobile digital microscopy using deep learning for the detection of soil-transmitted helminthes and *Schistosoma haematobium* eggs in human excreta, which could aid in medical diagnosis of infected people, but cannot improve identification or mapping of high risk environments. Here, we offer a new tool for automated detection and classification of larval parasites released from medically relevant freshwater snails in Senegal. The goal of this study was to provide a proof of concept that the application of the latest computer vision classification algorithms, powered by artificial intelligence (AI) technology, may help to improve snail and parasites identification in capacity-limited, resource-poor areas where schistosomiasis is endemic.

## Materials and Methods

### Datasets

During environmental monitoring of snails and parasites in the Senegal River Basin during 2016-2018, we collected 5,543 images of four genera of medically important and common freshwater snails -- (1) *Bulinus*, (2) *Biomphalaria*, (3) *Lymnaea*, and (4) *Melanoides* -- and 5,140 images of parasitic cercariae, from 11 morphospecies liberated from the *Biomphalaria* and *Bulinus* spp. snails encountered and dissected. **Table 1** summarizes the numbers of images in each category and **Fig 2** shows examples for each snail and parasite category in our dataset. Snail photographs were taken by mobile phone devices and cameras with similar resolution (at least 1024×768 pixels) and taken from similar angles, zoom, backgrounds, and lighting; while parasite images were taken from the same cell phones through dissecting microscopes, with the same uniformity. In addition, all parasite images were processed to grayscale. The removal of color was performed to further decrease lighting and time of day artifacts that could affect computer vision results. The detailed protocols for image collection and quality control are provided in the supplemental materials (see **S1 Appendix**). Initial identification of snails and parasites was performed by trained parasitologists in the field. Cases of disagreement between specialists were resolved.

**Table 1.** Summary of numbers of images for each snail and parasite species; numbers of images for the split of training (80%) and validation set (20%).

| Snail class | *Biomphalaria* | *Bulinus* | *Lymnaea* | *Melanoides* |
| --- | --- | --- | --- | --- |
| Number of images | 466 | 4215 | 725 | 137 |
| Training | 374 | 3372 | 580 | 109 |
| Validation | 92 | 843 | 145 | 28 |

| Parasite class | HS | NHS1 | NHS2 | AM | BO | EC | GY | ME | PP | PT | XI |
| --- | --- | --- | --- | --- | --- | --- | --- | --- | --- | --- | --- |
| Number of images | 1008 | 638 | 107 | 442 | 224 | 332 | 152 | 196 | 806 | 231 | 1004 |
| Training | 806 | 510 | 86 | 354 | 179 | 266 | 121 | 156 | 645 | 185 | 803 |
| Validation | 202 | 128 | 21 | 88 | 45 | 66 | 31 | 40 | 161 | 46 | 201 |

Abbreviations of parasite classes as follow, HS: Human-schisto, NHS1: Nonhuman- schisto forktail type I, NHS2: Nonhuman- schisto forktail type II, AM: Amphistome, BO: Bovis, EC: Echino, GY: Gymno, ME: Metacerc, PP: Parapleurolophocercous, PT: Parthenitae, XI- Xiphidiocercariae.

**Figure 2.**
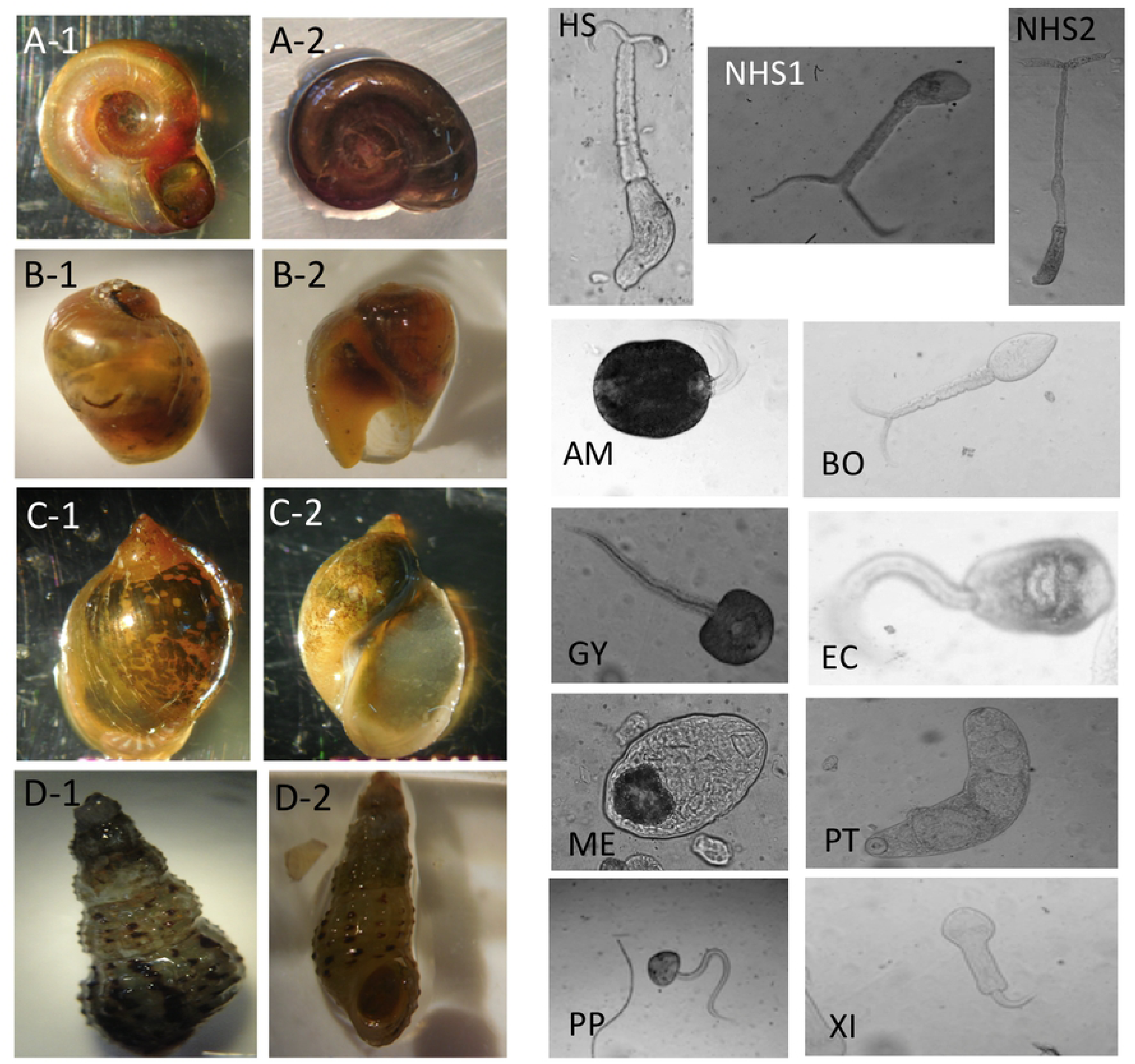
Image examples for snail and parasite classes. For snail classes, A-1 and A-2: Biomphalaria. B-1 and B-2: Bulinus. C-1 and C-2: Lymnaea. D-1 and D-2: Melanoides. For parasite classes, HS: Human-schisto, NHS1: Nonhuman-schisto forktail type I, NHS2: Nonhuman-schisto forktail type II, AM: Amphistome, BO: Bovis, EC: Echino, GY: Gymno, ME: Metacerc, PP: Parapleurolophocercous, PT: Parthenitae, XI-Xiphidiocercariae.

Accuracy of field identifications were verified by molecular barcoding technique: at the time of shedding or dissection, in all fork-tailed cercariae liberated from snails were placed individually on WhatmanFTA^©^ cards and sequenced to distinguish human-infective *Schistosoma haematobium* and *S. haematobium–bovis* hybrids from *S. bovis* (schistosome parasite of cattle) and other non-human furcocercous (forktailed) trematode species [8]. The identification of the parasites was based on multi-locus analyses with one mitochondrial (*cox*1) and two nuclear (*ITS*1+2 and 18S) genes [9]. Cercariae on FTA cards were accessioned into the Schistosomiasis Collection at the Natural History Museum (SCAN) [10].

All *Biomphalaria* and *Bulinus* spp. snails were also identified to morphospecies by both trained parasitologists and genetic barcoding. Total genomic DNA was isolated from a small amount of snail tissue using the DNeasy Blood and Tissue kit (Qiagen, UK) according to manufacturer’s instructions. Amplification of a partial cytochrome oxidase 1 (*cox1*) sequence was carried out on snail vouchers [11]. PCR and sequencing conditions were chosen as previously published [12]. Sequencing was performed on an Applied Biosystems 3730XL analyser (Life Technologies, UK) [9]. The field ID guidelines for morphologies of snails and parasites are provided in the supplemental materials (see **S2 Appendix**).

For imagery of snails and parasites, we strategically avoided extensive processing of images, since the ultimate goal is to create a CNN capable of classification of images of variable quality and resolution. We anticipate that many technicians will be taking pictures directly from mobile devices in the field. So, in the preparation of our training data, images that were severely blurry were removed from the test and validation sets, but were still used in training. For each class of snails and parasites, we randomly divided the dataset into training and validation set, containing 80% and 20% of the total images, respectively (**Table 1**). There was no overlap between the training set and the validation set.

### Training Algorithm

Deep learning algorithms with recent advances in computation and large datasets have been shown to be comparable with, and even exceed, human performance in various object recognition and computer vision tasks, including applications to diagnose human disease (e.g., ImageNet challenge [13], breast cancer histology images [14], and skin cancer images [15]). Convolutional neural networks (CNNs) learn key features directly from the training images by the optimization of the classification loss function [16,17] and therefore have minimal need for *a priori* knowledge to design a classification system. Thus, the performance is less biased by the assumptions of the researchers [13,18].

In this study, we utilized VGG16 pre-trained model [19], a 16-layer CNN architecture, which was trained on approximately 1.28 million images with 1,000 categories. VGG16 was used by the VGG team (Visual Geometry group at the University of Oxford) in the 2014 ImageNet Large Scale Visual Recognition Challenge. We trained VGG16 on our snail and parasite images using transfer learning [20]. Transfer learning is defined as exporting knowledge from previously learned sources to a target task [21]. The network architecture established here has convolutional-pooling layer pairs (max-pooling), followed by a fully-connected network [17]. The network architecture is illustrated in **S1 Fig**. The CNN is trained end-to-end directly from image labels and raw pixels, with a VGG16 network for photographic images of snails and another separated network for microscopic images of parasites.

We adopted the training patches as 64×64 pixels for input layers in the consistent with VGG16 network architecture, to ensure the patch size was sufficient to cover the relevant structures and morphologies of snail and parasites. We first initialized the weights with the VGG-16 pre-trained on ImageNet dataset, then froze the bottom of the network and just trained the top of the VGG-16 convolutional networks (**S1 Fig**). The fully-connected layers were composed of Rectified Linear Units (i.e., the ReLU activation function), to avoid the vanishing gradients and to improve the training speed [22]. The output layer was composed of 4 neurons for snail classification, corresponding to each of the four classes that are normalized with a softmax activation function. For the parasite classification task, 11 neurons were set up in the same manner. The model was trained with 80% of the training set, and validated on the remaining images not used for training. Note that the validation set is randomly selected for each epoch, which is the measure of the number of times all of the training images are used once to update the network weights [17]. The network weights were initialized randomly, and an adaptive learning rate gradient-descent back-propagation algorithm [22] was used for weight update. Here we selected categorical cross-entropy as a loss function in the model. In these two classification tasks, the CNN outputs a probability distribution over 4 classes of snails and 11 classes of parasites.

Since our dataset was rather small comparing with recent studies using CNNs for image recognition tasks [13], here we applied the techniques of dropout and regularization to overcome the overfitting of training data [17]. We implemented the CNN model with VGG16 network in a Python environment with the Keras package [23], and Google’s deep learning framework, TensorFlow [24].

### Data Augmentation

CNN algorithms work best for balanced dataset, i.e., when the categories of objects that need to be identified are equally represented in the dataset [17]. Our dataset was unbalanced: for example, *Bulinus* snails comprised about 75% of total snail images, since this snail genus was most common in the field sites. Thus, we applied a data augmentation approach and used *rotation* and *shifting* [25] to generate more images for the snail and parasites species that were underrepresented in the dataset. *Mirroring* was not used *Bulinus* and *Lymnaea* spp due to the diagnostic value of their coil orientation (i.e. *Bulinus* is sinistral and Lymnaea is dextral). We also applied Gaussian noise to the background of parasite training images [26] to ensure that the model did not learn background artifacts, but focused only on learning the morphologies of the parasite objects. In this study, we used a Keras defined generator for automating data augmentation [23]; every item of every batch was randomly changed according to the following settings: (1) rotation range = 20, (2) width shift range = 0.2, (3) height shift range = 0.2, and (4) shear range = 0.15. In practice, rotations and shifting allowed us to increase the size of the dataset without deteriorating its quality. Data augmentation used here further improved the datasets and CNN predicting performance.

### Model deployment

After building and training the CNN model, we then deployed the trained model and network weights to establish a web application for inference using TensorFlow.js [27], which is a library for executing machine learning algorithms in JavaScript. TensorFlow.js is compatible with the Python-based TensorFlow and Keras APIs, allowing our Keras model to be converted to a JavaScript format that can be run in a web browser. This makes it accessible on any device with a modern browser including both smartphones and common laptop computers. Using browser storage and caching APIs, the web application can even be used in areas with limited or no internet connectivity.

### Ethics Statement

Freshwater snails were collected in collaboration with Centre de Recherche Biomédicale Espoir pour la Santé in Senegal who obtained the permission to conduct the field snail collection, from The Direction de l’Environement et des Etablissements Classés with the identification number “N°002302 MEDD/DEEC/yn”. In this study, we use photos of snails for the machine learning model training and validation, therefore, there was no direct use of snails in the field environment.

## Results

We evaluated our CNN model’s performance with metrics of accuracy, sensitivity, specificity, and F1 score on the test set. With the optimized CNN architecture, we obtained 99.6% accuracy (proportion of correct classifications, either true positive or true negative) for the 4 snail genera and 88.14% for the 11 parasite morphospecies. The optimized dropout rates for the convolutional layer [17] for the snail dataset was 0.6 and for parasite dataset was 0.75. For the parasite set, we ran a second analysis with only 3 classes of parasites relevant to map risk for schistosomiasis transmission, namely: human schistosomes, non-human forktail cercariae, and other trematode morphotypes. The overall accuracy for the 3 parasite classes increased to 92.41%. The sensitivity of distinguishing human schistosomes from non-human parasites was 83.66%. **Fig 3** shows the confusion matrix of our method over the 4 snail general and 11 parasite morphotypes, along with details of true positive (TP), true negative (TN), false positive (FP), and false negative (FN) outcomes.

**Figure 3:**
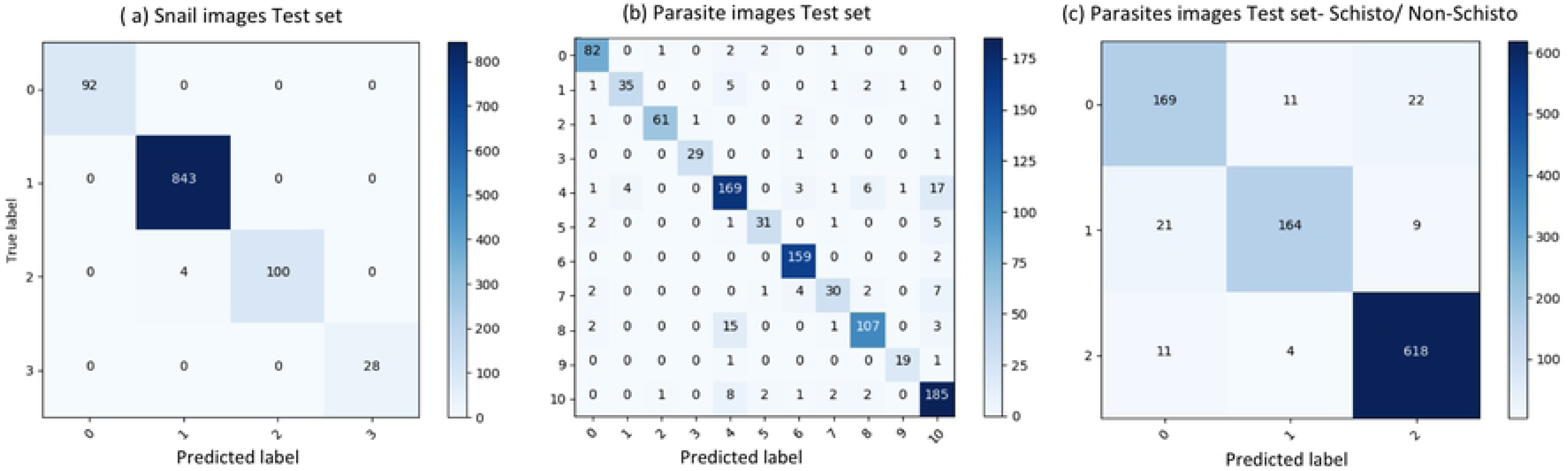
Confusion matrix of classification on test set. (a) results of snail image set, labels: 0-Biomphalaria, 1: Bulinus, 2: Lymnaea, 3: Melanoides. (b) results of parasite image set, labels: 0: Amphistome, 1: Bovis, 2: Echino, 3: Gymno, 4: Human-schisto, 5: Metacerc, 6: Parapleurolophocercous, 7: Parthenitae, 8: Nonhuman-schisto forktail type I, 9: Nonhuman-schisto forktail type II, 10: Xiphidiocercariae. (c) Combining other trematodes as one category, labels: 0: Schisto, 1: Non-human forktail type I, type II, and Bovis, 2: Other trematodes.

Sensitivity, specificity, recall, precision, and F1 score for each categories were calculated as follows:

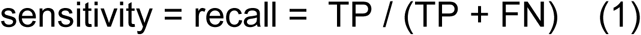

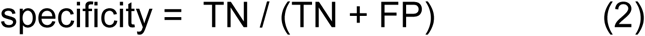

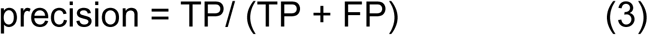

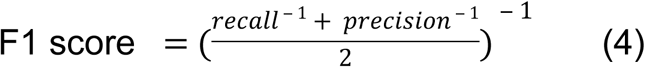

Sensitivity measures the proportion of positives that are correctly identified; in our case, we focused on the percentage of human schistosomes correctly identified. Specificity measures the proportion of negatives that are correctly identified; in our case, we focused on the percentage of non-human forktailed cercariae (some of which can potentially be visually similar and hard to distinguish from human schistosomes) that are correctly identified as non-human schistosomes. Precision is a measure of a classifier’s exactness, while recall is a measure of a classifier’s completeness. Low precision indicates a large number of false positives, while low recall indicates many false negatives. F1 score conveys the balance between precision and recall and is defined as the harmonic mean of precision and sensitivity. Our results showed that the CNN produced high sensitivity and specificity (> 84%) as well as F1 score (> 0.84) in all categories. The metrics to measure classification performances are shown in **Table 2** and demonstrate the robustness of the CNN training algorithm for the tasks of image recognition for both snails and parasites, despite the low sample size we had to work with.

**Table 2.** Results of sensitivity(%) and specificity(%) of CNN model.TP: true positive, TN: true negative, FP: false positive, FN: false negative.

| Snail class | TP | TN | FP | FN | sensitivity(%) | specificity(%) | F1 score |
| --- | --- | --- | --- | --- | --- | --- | --- |
| Biomphalaria | 92 | 1020 | 0 | 0 | 100.00 | 100.00 | 1.00 |
| Bulinus | 843 | 265 | 4 | 0 | 100.00 | 98.51 | 0.99 |
| Lymnaea | 145 | 963 | 0 | 4 | 97.31 | 100.00 | 0.99 |
| Melanoides | 28 | 1084 | 0 | 0 | 100.00 | 100.00 | 1.00 |

| Parasite class | TP | TN | FP | FN | sensitivity(%) | specificity(%) | F1 score |
| --- | --- | --- | --- | --- | --- | --- | --- |
| Schisto | 169 | 795 | 32 | 33 | 83.66 | 96.13 | 0.84 |
| Non-human forktail | 164 | 820 | 15 | 30 | 84.53 | 99.51 | 0.91 |
| Other trematodes | 618 | 365 | 31 | 15 | 97.63 | 92.17 | 0.96 |

### Comparison with human parasitologist performance

To validate our deep learning approach, we compared the direct performance of the CNN to 8 human parasitology experts. For each image, the parasitologists were asked to identify the category of the snails and parasites. We prepared 30 snail images from among the 4 categories and 120 parasite images from among the 11 morphotypes in the computer’s validation sets. For each test, previously unseen, molecularly verified images of trematode cercariae and snails were displayed, and parasitologists are asked to identify them from among the same categories of snails and parasites that the computer vision algorithm trained on. Identification guidelines were provided along with the quiz. The sample quiz is included in supplement materials (see **S3 Appendix**). The metrics to measure human parasitologists’ performances and compare with the computer’s, such as sensitivity, specificity, rand F1 score are shown in **Table 3** and **Fig 4**. The CNN generated a malignancy probability *P* per image. We then fix a threshold probability *t* such that the prediction *ŷ* for any image is *ŷ = P ≥ t*, and the blue curve (**Fig 4**) is drawn by sweeping t in the interval 0–1. The area under the curve (AUC) is the CNN’s measure of performance, with a maximum value of 1. The AUC for human schistosomes, non-human forktailed cercariae, and other trematodes were 0.96, 0.95, 0.98, respectively. The CNN achieved superior performance to a parasitologist if the sensitivity– specificity point of the parasitologist lies below the blue curve, which most do. Note that sensitivity and specificity of human parasitologists to correctly distinguish cryptic human schistosome and non-human-schistosome species (*Schistosoma bovis)* was significantly lower than the CNN.

**Table 3.**
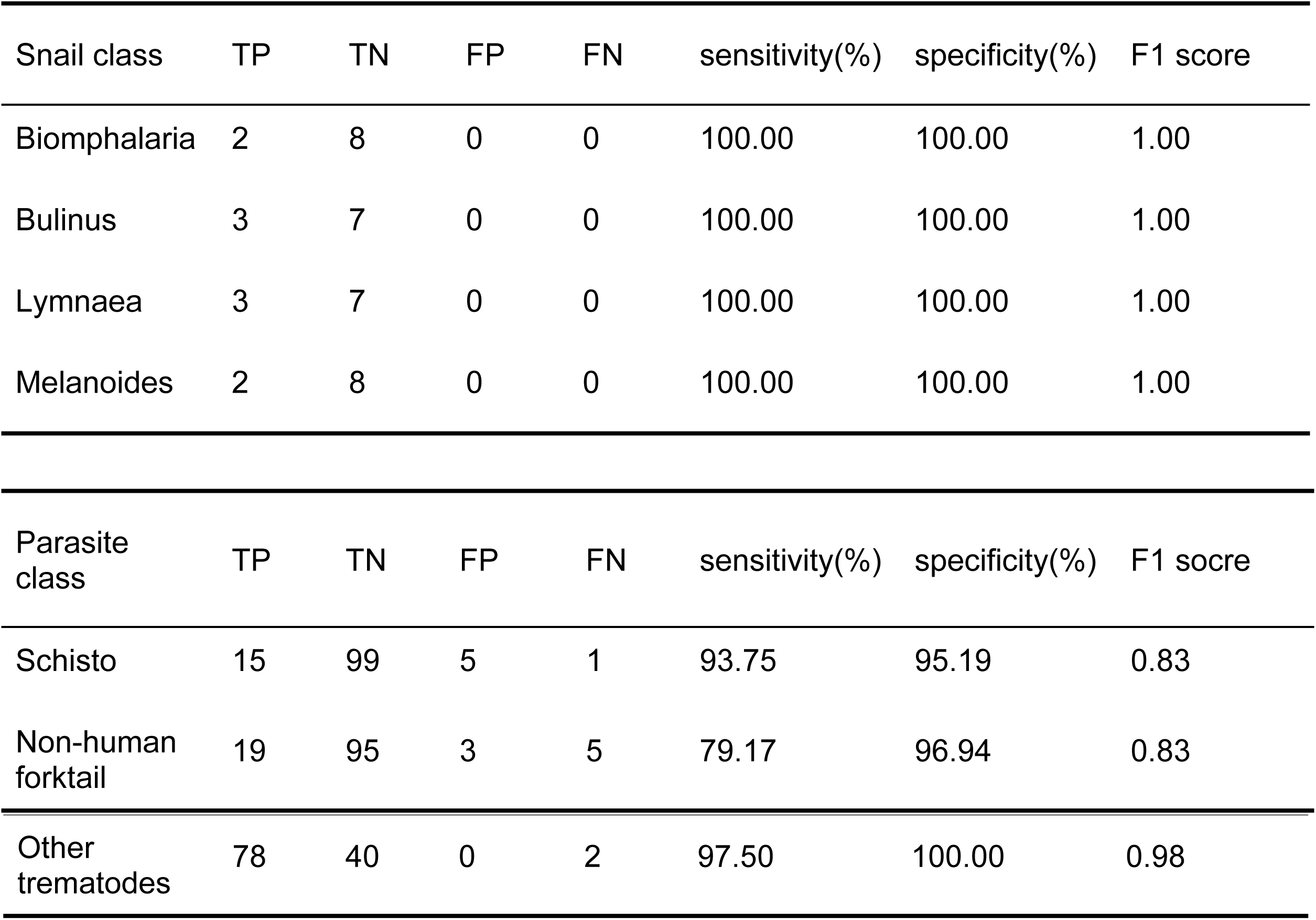
Results of sensitivity(%) and specificity(%) of 8 human parasitologists

**Figure 4:**
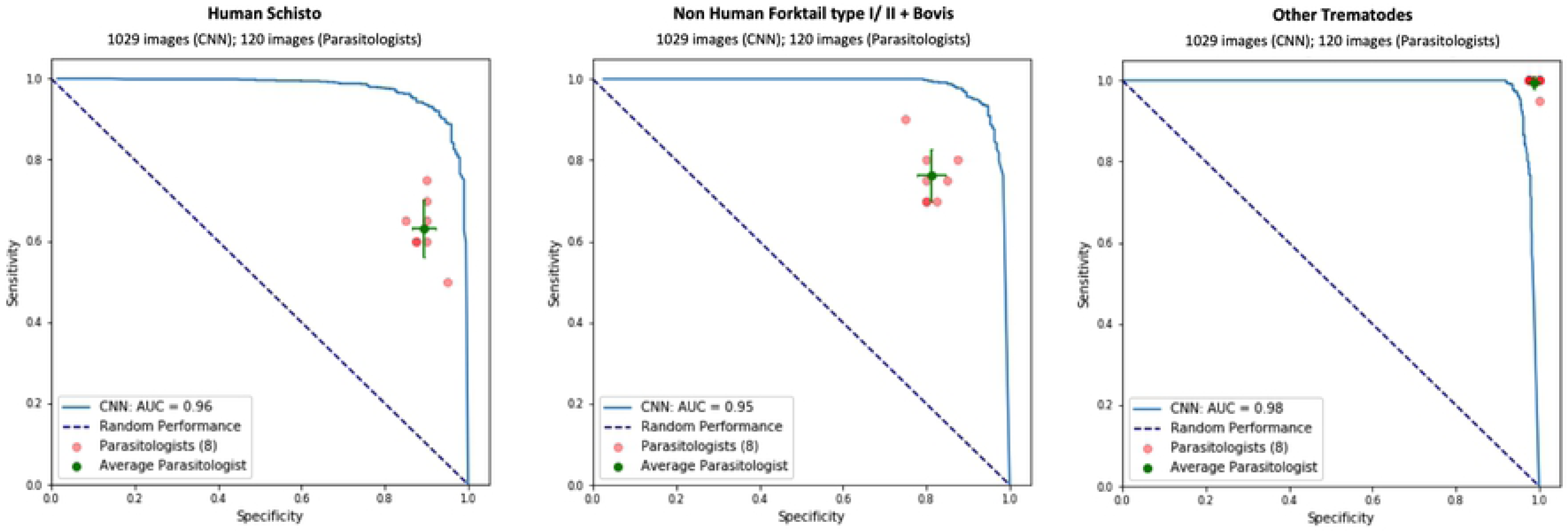
Comparison of classification performance with CNN and parasitologist. The CNN outperforms any parasitologist whose sensitivity and specificity point (in red color) falls below the blue curve of the CNN. The green points represent the average of the parasitologists (average sensitivity and specificity of all red points), with error bars denoting one standard deviation. We simplify the 11 classes of parasite to only 3 classes of interest for schistosomiasis environmental risk mapping: (a) human schistosomes, (b) non-human forktail cercariae, and (c) other trematode morphotypes. The area under the curve (AUC) for each case is over 95%. The CNN achieves superior performance to a parasitologist if the sensitivity–specificity point of the parasitologist lies below the blue curve.

## Discussion

In this work we developed and presented a deep learning model that is capable of accurately classifying images of human Schistosomiasis larval parasites and intermediate host snails to benefit environmental monitoring and control of this debilitating disease. The accuracy of our models matched and in most cases exceeded highly trained field experts that we tested it against. Our convolutional neural network (CNN) classification performed better for snail recognition than for parasite recognition. This is because snail morphologies are generally more distinct, and it is easier to take high quality pictures of snails than parasites. This decreases the risk of erroneous classification of snails, as reflected by the small number of false positives and false negatives in our model evaluation. Parasites, on the other hand, show more complex morphologies and variation, even within a species, which makes discrimination of trematode cercariae more challenging, as reflected by the slightly large number of false negative and false positive classification.

A benefit of CNN is that models can be trained with rather limited image datasets using data augmentation approaches. Because we detected a low number of morphologically similar forked-tail human and non-human schistosoma parasites, we used rotation and shifting methods to effectively increase the number of images used in training. We had the advantage of also having video footage of living, mobile cercariae. As in our case, in the future, collecting video imagery of parasites to generate many unique still-images of cercariaes an easy way to increase samples for training CNN models.

In addition to building a deep learning model, we have also developed a workflow for image collection and quality control for other researchers to follow, to quickly build an image database that can facilitate identification of medically-important snails and their parasites with the click of a cell phone and a pre-trained computer vision model. The protocol for image collection is provided in the supplemental materials (see **Appendix S2**). We hope that user photos can ultimately be used to create a global database of medically important snail and parasite species, and the species they co-exist with that are not relevant to human health, useful for further model training. When collecting the samples in the field, we also suggest that GPS locations be recorded, which can facilitate the application of these data for precision mapping of snail and parasite distribution in space and time. We take inspiration from online platforms that encourage citizen science and crowdsourcing activities, similar to iNaturalist, NaturNet, MycoMap, etc. iNaturalist, for example, has the image classification capacity to immediately suggest an object name right after a user has uploaded an image to the platform.

To encourage the development of our specific platform and for translating it for use on other medically-relevant parasites, vectors, and non-human hosts found in the environment, our code, image sets, and neural network weights are provided in a public repository, which also includes detailed documentation to assist researchers in following our workflow to (1) reproduce the results, and (2) build new models using their own image datasets. We have also deployed our classification model to a web application that allows researchers to select their own images of snail and parasites to obtain the classification prediction, which is the probability of an object of belonging to a specific class. Even though it takes a considerable computing power to properly train a CNN image classification model, once trained and deployed, a CNN can provide a result in a fraction of a second on a smartphone and a common laptop computer.

In conclusion, this study demonstrates the effectiveness of deep learning in image recognition tasks for classification of medically relevant snails and their parasitic infections. We apply a computer vision model, using a single convolutional neural network trained on a relatively low number of cell-phone and dissecting microscope acquired images. Thus, we have developed a proof of concept for this technology to be applied in resource-poor settings where schistosomiasis is endemic and where identification of hotspots of transmission is desperately needed to target interventions. Our model matched the performance of 8 highly trained human parasitologists familiar with snail and parasite diversity in the study region. In light of its success, we’ve deployed our product as a publically-accessible web application. In the future this method could be deployed on mobile devices with minimal cost, and holds the potential for substantial improvement for monitoring and identifying snail and schistosomiasis hotspots. Deep learning is a powerful tool that can help fill the capacity gap limiting our understanding of the environmental components of transmission for more affordable, efficient, and effective control of neglected tropical diseases.

## Acknowledgement

This work was supported by a grant from the Bill and Melinda Gates Foundation (OPP1114050), an Environmental Ventures Program grant from the Stanford University Woods Institute for the Environment, a SEED grant from the Freeman Spogli Institute at Stanford University, a Human Centered Artificial Intelligence grant from Stanford University, a grant from the National Institutes of Health (R01TW010286), and a grant from the National Science Foundation (1414102). We thank Tim White and Richard Grewelle for their suggestions and comments for this project. We thank the parasitologists who participated in the quiz that was designed to compare the performances of CNN and human experts on snails and parasites image classification. We also thank IBM open source group, especially Yi Hong Wang, Chin Huang, Ted Chang, Catherine Diep, Winnie Tsang, and Thomas Truong for providing guidance, education, and cloud resources to establish the CNN model deployment process and host the web application. Open Source code packages, image dataset, and web application can be accessed via this link (https://github.com/deleo-lab/schisto-parasite-classification).

## Author Contributions

ZYCL and AC built, trained, and validated the CNN model.

AC, IJJ, and SHS organized and labeled the image dataset.

AC, IJJ, SHS, GR, NJ, RN, and GADL conducted the field works.

PS performed the initial training and testing.

PVE and TN deployed the CNN model and created the web application.

GADL conceived the idea and supervised the project in all perspectives.

## Supporting Information Legends

### Supplement Figure S1

VGG-16 architecture illustration.

### Supplement Appendix

S1. Protocols for image collection and quality control

S2. Field ID guidelines for morphologies of snails and parasites

S3. Sample quiz on snail and parasite classification for human parasitologists

